# Development of lyophilized faecal inoculation capsules for use in koala rehabilitation and conservation

**DOI:** 10.64898/2026.09.28.754943

**Authors:** Michaela D.J. Blyton, Leanne Dierens, Max Lacour, Philip Hugenholtz

## Abstract

As a specialist herbivore, koalas rely on their gut microbiomes to help digest their toxic and fibrous diet of *Eucalyptus* leaves. Without these critical microbes, koalas may not be able to obtain the nutrients and energy they need to survive. As such, a large proportion of koalas that undergo rehabilitation for chlamydiosis develop gut dysbiosis from the antibiotic treatment and are either euthanized or die. Here we aimed to modify previously developed fresh faecal inoculation capsules for an extended shelf-life such that they could be applied in a clinical setting to prevent or treat gut dysbiosis. Using 16S rRNA gene amplicon sequencing we demonstrated that air-drying faecal material leads to an overgrowth of facultative anaerobes, whereas lyophilised material retains a similar microbial composition to fresh material. Initial survival of koala faecal microbes as assessed by live/dead staining combined with microscopy was high after lyophilisation regardless of which excipient was used, except for 20% glycerol that resulted in ∼15% lower survival than other treatments. Lyophilised fine particles extracted from koala faeces and packaged into acid-resistant capsules maintained their original microbial composition and had high microbial survival over a year, regardless of the excipient used. Dry-fill, single-layer capsules maintained integrity after 10 hrs in synthetic koala stomach acid. These capsules may be useful in adapting the gut microbiomes of koalas to novel diets e.g. during translocations. However, attempts to apply the capsules in the clinical setting were unsuccessful due to unanticipated difficulties in administrating the capsules to sick koalas.

**Importance:** In recent years wildlife conservation has begun to appreciate the importance of the gut microbiome to animal nutrition, health and fitness. This has led to the increasing use of probiotics and faecal transplants to change or support the gut microbiomes of animals during translocation, reintroduction and rehabilitation. Unfortunately, in many instances it is difficult to source appropriate inoculant for these interventions and what is available often has limited viability. In this study, we developed and tested acid-resistant faecal inoculation capsules for use in koalas and showed that they have a shelf-life of at least a year at room temperature. These capsules may be useful in adapting the gut microbiomes of koalas to novel diets during translocations and the lyophilisation approach will contribute to the ongoing development of strategies to prevent gut dysbiosis during antibiotic treatment. The design of the capsules could also be adapted for use in other wildlife species.

## Introduction

Northern koala populations in Australia continue to decline due to habitat loss and disease^1,2^, in particular, the 2020 mega-fires devastated previous koala strongholds leading to koalas being listed as endangered in the Australian states of Queensland, New South Wales and the Australian Capital Territory in 2022^3^. As such, it is now more critical than ever that sick and injured koalas brought into rehabilitation centres^4^ can be returned to health and released so that they can contribute to the ongoing viability of local populations. Approximately half of koalas admitted to wildlife hospitals are given antibiotics to treat chlamydial disease that causes cystitis, reproductive pathology and/or conjunctivitis, often progressing to infertility and/or mortality^2,5^. These antibiotic treatments can decimate the koalas’ community of symbiotic gut microbes (microbiome) causing gastrointestinal dysbiosis that can contribute to mortality^6^, with over 60% of koalas treated for disease being euthanised^5^. As specialist herbivores, koalas rely on their gut microbiomes to help digest their toxic and fibrous diet of *Eucalyptus* leaves^7,8^. Additionally, in other species including humans the gut microbiome has been found to prevent infection by gastrointestinal pathogens^9^. Without these critical microbes, koalas may not be able to obtain the nutrients and energy they need to survive, leading to poor rehabilitation outcomes for antibiotic-treated koalas^5,10^. In this project, we sought to modify and optimise fresh faecal inoculation capsules that were previously developed for research purposes, into a product with an extended shelf-life that could be used in a clinical setting to treat or prevent gut dysbiosis in koalas that receive antibiotics.

Sick koalas and those that die during antibiotic treatment have gut microbiomes that differ from those of healthy individuals, with koalas that recover from antibiotic treatment regaining a microbiome community similar to that of healthy koalas^10–12^. This suggests a link between koala health and their symbiotic microbes. Koalas have a greatly enlarged caecum and proximal colon to accommodate the microbial fermentation of dietary fibre (hemicellulose and cellulose) into short chain fatty acids that koalas use for energy^13^. The koala gut microbiome has been shown to vary between individuals that habitually feed on different eucalypts^14,15^ and between koala populations^16^ but the overall community does not appear to respond to diet change^17^. Rather, we have found that the gut microbiome may influence which species of eucalypts individual koalas can feed on^14,17^. In turn this could constrain habitat selection and limit the survival of translocated or rehabilitated koalas after release. Therefore, maintaining an appropriate gut microbiome throughout antibiotic treatment could not only improve survival outcomes in rehabilitation by maintaining digestive efficiency and preventing dysbiosis but could also have longer term benefits when those koalas are returned to the wild.

Wildlife carers and veterinarians have long realized the critical importance of maintaining the gut microbiomes of injured and sick koalas during antibiotic treatment. Yet, few suitable options are available to them. Some veterinarians utilize pre and probiotic formulations developed for humans, domestic animals or livestock that are inappropriate for use in koalas due to their distinct gastro-physiology and diet. Others use ‘poo shakes’ composed of ground koala faeces from putatively healthy koalas. With no guidelines for dose or preparation techniques by wildlife carers, these shakes often have low microbial density and diversity which lead to dubious benefits to the koalas. Currently, the best microbial inoculum for unhealthy koalas is caecal contents harvested from dead healthy koalas. However, few suitable koalas are available and the caecal contents cannot be stored for long periods before use. Further, it is unknown what proportion of the microbes contained in the shakes or caecal contents survive transit through the highly acid koala stomach^8,18^.

As part of our research into the koala gut microbiome, we previously developed microbial inoculation capsules that can be easily administered orally to wild or captive koalas under conscious restraint using a “pill popper”^14^. Through laboratory testing, we designed a capsule that can protect faecally derived microbes during their 10-hour journey through the acidic environment of a koala’s stomach. The microbes are then released in the neutral pH environment of the koala’s small intestine. We have also shown that when koalas are administered with two capsules daily for nine days, the introduced microbes are able to establish and persist beyond the completion of the treatment^14^. Thus, these capsules have the potential to stimulate the growth of beneficial microbes to support and restore koala gut health, akin to the use of probiotics in human medicine.

In our original design, the capsules contained live microbes concentrated from fresh faeces that had to be administered to koalas within a few hours of preparation. The presence of liquid in the inoculum internally weakened the capsules, which are designed for dry fill. This made the capsule assembly process intricate and time consuming, while also limiting the capsules to immediate use. Additionally, it is unknown how long the microbes can survive outside the anaerobic environment of the gut inside the capsules. In this study we resolve this issue by developing a dry-fill variant of the capsules. Lyophilising (freeze-drying) microbes before they are packaged into capsules is a commonly used technique in the human pharmaceutical industry to preserve probiotic bacteria and prolong shelf life^19–22^. However, not all microbes can survive this process and while excipients can improve survival it is unclear whether they can maintain a representative microbial community or bias the surviving microbes towards particular taxa^19^. Further, freeze driers are not readily available in a wildlife veterinary setting. Therefore, here we assessed: 1) the initial survival of koala faecal microbes and viable cell taxonomic composition after air-drying or lyophilisation with a range of different excipients; 2) the survival of koala faecal microbes and viable cell taxonomic composition over the course of a year at room temperature after lyophilisation and packaging into acid-resistant capsules; and 3) the optimal design of the acid-resistant capsules for dry-fill formulations.

## Results

### Assessing initial survival and composition of faecal microbiomes post preservation

In our first trial we tested the initial survival of microbes derived from freshly collected captive koala scats preserved using nine different treatments (Table 1). Both the air-dried ground whole scats and the air-dried fine particles had high viable cell proportions (49.5 and 30.8%, respectively) of facultative anerobic bacteria belonging to the family Enterobacteriaceae (including *E. coli* and the genera *Enterobacter* and *Klebsiella*), while Enterobacteriaceae only had a relative abundance of 0.1% in the fresh material (Figure 1a). Further, the facultative anaerobes belonging to the family Morganellaceae were only detected in the air-dried treatments suggesting that this treatment favours survival of facultative anaerobes and may even allow for their growth during drying (Figure 1a).

**Figure 1:**
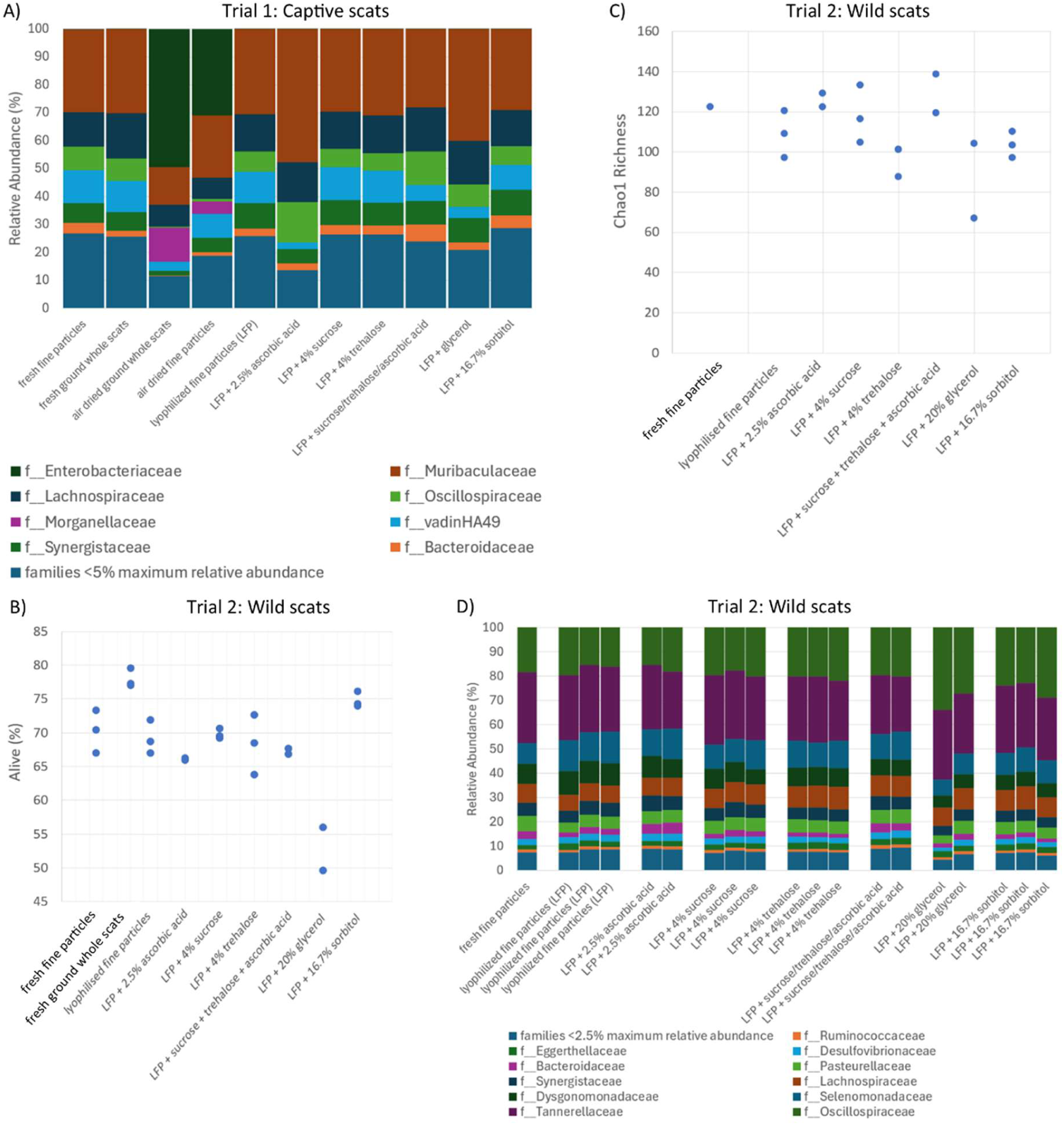
A) Bar plot showing the viable cell relative abundances of the major (>5% max within sample relative abundance) families in the first trial of initial preservation of koala faecal microbes from captive koala scats across treatments. B) Proportion of cells that were initially viable after presentation for each treatment in the second trial which used wild koala scats as the starting material. C) Viable cell richness measured by Chao1 Index from the feature counts across the treatments in the second trial. D) Bar plot showing the viable cell relative abundance of the major (>2.5% max within sample relative abundance) families in the second trial.

**Table 1:** Preservation formulations assessed for initial microbial survival and viable cell taxonomic composition.

| Formulation | Selected for Longevity study |
| --- | --- |
| 1. Fresh fine particles (Control) | N/A |
| 2. Fresh ground whole scats (Control) | N/A |
| 3. Air dried ground whole scats | No – enriched for facultative anaerobes |
| 4. Air dried fine particles | No – enriched for facultative anaerobes |
| 5. Lyophilized fine particles | Yes |
| 6. Lyophilized fine particles + ascorbic acid (2.5% w/w)<sup>19,20</sup> | Yes – enriched for Muribaculaceae, highest initial richness |
| 7. Lyophilized fine particles + sucrose (4% w/w)<sup>22</sup> | Yes |
| 8. Lyophilized fine particles + trehalose (4% w/w)<sup>22</sup> | No – lower initial richness |
| 9. Lyophilized fine particles + sucrose + trehalose + ascorbic acid* | No – multiple excipients without benefit |
| 10. Lyophilized fine particles + glycerol (20% v/w) | No – poorer initial survival, enriched for Muribaculaceae, lower initial richness |
| 11. Lyophilized fine particles + sorbitol (16.7% w/w)<sup>21</sup> | Yes – highest initial survival |

\* proportions as in formulations 6, 7 and 8

In the first trial, the family Muribaculaceae was also overrepresented in the viable cells from the lyophilised fine particles (LFP) + 2.5% ascorbic acid and LFP + 20% glycerol treatments compared to the fresh fine particles (47.6% and 40.0% vs 30.1%, respectively; Figure 1a). While this would suggest that those treatments do not preserve a representative koala faecal microbial community, Muribaculaceae is often overrepresented in captive relative to wild koala microbiomes^16^. Therefore, the experiment was repeated using material sourced from wild koalas (except for the air-dried treatments that were eliminated as viable preservation methods).

In the second trial, scats from wild koalas were used as the starting material. In that trial there were similar initial survival rates between most preservation treatments (average = 67-70%) and the fresh fine particles (average = 70%; Figure 1b). Apart from the LFP + 20% glycerol treatment, which had lower average microbial survival (53%) than the other treatments and the LFP + 16.7% sorbitol treatment that had higher average survival than the fresh fine particles (75%). Interestingly, survival was also higher in the fresh ground whole scats (78%) when compared to the fresh fine particles.

Fewer species survived preservation in the LFP + trehalose and the LFP + glycerol treatments when compared to the fresh fine particles, while LFP + 4% ascorbic acid had the highest species richness (as estimated from Chao1 diversity calculated from the 16S rRNA amplicon sequencing data; Figure 1c). All treatments had very similar viable cell compositions to that of the fresh fine particles (Bray-Curtis distance to fresh fine particles based on feature read counts: 0.085-0.151) except for one of the LFP +20% glycerol replicates that was more dissimilar (Bray-Curtis distance to fresh fine particles = 0.204; Figure 1d).

From these analyses, it was concluded that the four treatments with the best initial survival characteristics after preservation were lyophilised fine particles, LFP + ascorbic acid, FDFP + sucrose and FDFP + sorbitol based on the advantages and drawbacks of each treatment as summarised in Table 1.

### Assessing the survival and composition of preserved and encapsulated faecal microbes over a year

The four best performing treatments from the assessment of initial survival were assessed for how well the preserved microbes survived over the course of a year when packaged into hypromellose capsules (DRCaps, Capsugel®) coated in shellac and kept at room temperature in amber jars.

In the first three weeks after the capsules were assembled, the proportion of viable cells decreased across all preservation treatments from 69-72% to 58-59% (Figure 2a). After this time the proportion of viable cells stabilised and remained stable across treatments (with some fluctuations likely due to reconstitution variation) out to 7 months (217 days). Then between 7 and 9 months (287 days) the proportion of viable cells decreased across treatments to 50-51%. From 9 months to one year the proportion of viable cells remained stable at approximately 50% across all treatments.

**Figure 2:**
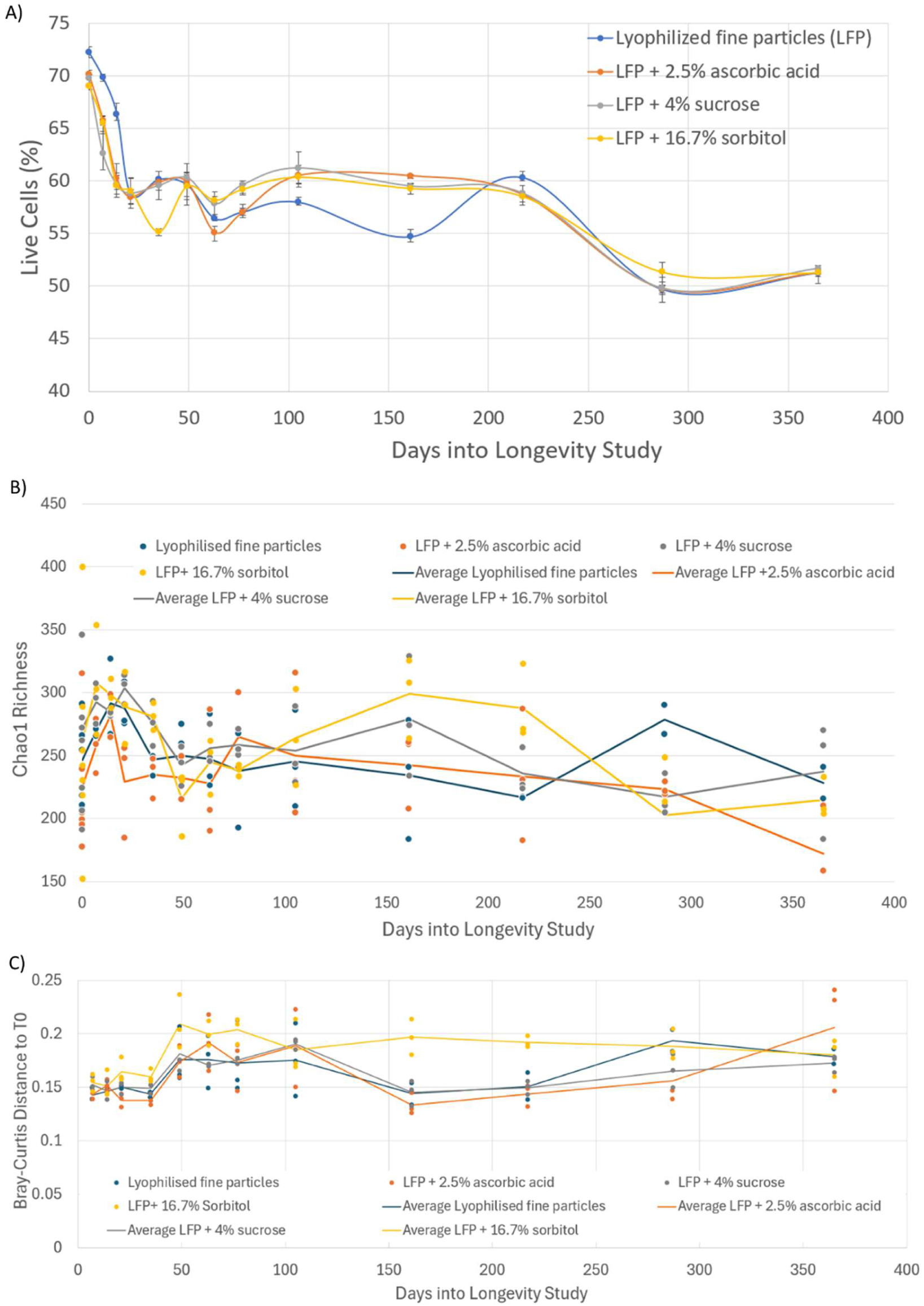
A) Proportion of cells that remained viable; B) Adjusted Chao1 richness; and C) Adjusted Bray-Curtis distances to the corresponding time zero samples over the course of a year after presentation and encapsulation for each treatment.

There was no significant linear change in richness over the course of the year for any of the treatments (Chao1 richness; p = 0.885; Figure 2b). However, across time points the LFP + 2.5% ascorbic acid treatment was found to have significantly lower richness compared with the other treatments (pairwise t-tests: 0.002 < p < 0.038; Figure 2b). Overall, there was no significant linear relationship between the Bray-Curtis distance of a sample to the time zero sample and the number of days since the capsules were prepared for any of the treatments (treatment x day: p = 0.275; day: p = 0.7721; Figure 2c). Although, on inspection of the adjusted Bray-Curtis distances it was observed that dissimilarity to the time zero samples was generally lower for the first sequencing run (days 7-35; Figure 2c). This may be explained by the fact that the time zero samples and the samples on the first sequencing run were extracted in the same batch leading to lower dissimilarities. Across the course of the year, the LFP + 16.7% sorbitol treatment had significantly high dissimilarity to the time zero sample when compared with the dissimilarities of the other treatments to their time zero samples (p < 0.001), suggesting that there was a greater shift in community structure after preservation for the sorbitol treatment. However, given that it did not increase over time, the change must have occurred in the first few days after preservation and then remained stable. Therefore, in all treatments there was little change in viable cell community composition over the course of the year.

### Determination of optimal capsule design for dry-fill

The *in vitro* experiment of capsule integrity showed that both the double and single layered hypromellose capsules (DRCaps, Capsugel®) coated in shellac, maintained integrity in an acid solution (typical of the koala stomach) for at least 10 hours and broke down in an alkaline solution (typical of the koala hindgut) within an hour as indicated by the discolouration of the alkaline solution by crystal violet when released from the capsules.

## Discussion

### Successful development of koala faecal inoculation capsules with prolonged shelf-life

This study aimed to refine the design of previously developed koala faecal inoculation capsules such that they could be stored at room temperature for an extended period to facilitate their use in a rehabilitation setting. Through a series of experiments we identified that lyophilised fine particles extracted from koala faecal material and packaged into a single layer acid resistant capsule produced a product that i) maintained bacterial survival and a representative microbial community composition for at least 12 months; ii) may avoid currently unknown side effects in recipient koalas produced by including excipients; and iii) should maintain structural stability through transit of the koala stomach (as indicated by an *in vitro* trial), while permitting an increased dose of inoculum relative to the original double layered capsules.

The initial preservation experiment suggested that some cell death does occur during extraction of the fine particles from the whole scats. However, in our previous work we have shown that fine particles have between 200-300% greater cell densities compared with whole scats^14^. Thus, this increased cell density more than offsets the 8% decrease in survival making fine particles superior to whole ground scats for inoculation.

The initial decrease in survival observed across all preservation treatments during the first three weeks of the longevity study was most likely due to the death of cells that were not well persevered initially. The reason for the second decrease between 7 and 9 months is unknown but was not associated with the death of particular taxa and as such a viable cell community representative of the starting faecal material was maintained across the year. Further, over 70% of the initially viable cells remained alive over the course of the year across all treatments.

### Optimal preservation methods for the koala faecal microbiome

Our first trial of initial microbial preservation revealed that air-drying scats is not a recommended preservation method for the koala microbiome as it can result in overgrowth of facultative anaerobes that could potentially include opportunist pathogens such as *E. coli*^36,37^. This is in contrast to a study of koala scat aging that did not find an increased relative abundance of facultative anaerobes in older scats^38^. However, this may be because the DNA in that study was extracted from the centre of the faecal pellets, which is less exposed to oxygen, while in this study the whole scat was processed post drying. Alternatively, it may be that the growth of facultative anaerobes is variable based on the microbial composition of the scat on defecation and the climatic condition during drying/aging. In either case, the production of faecal inoculum in rehabilitation clinics that do not have access to freeze-drying equipment is not recommended due to the potential for overgrowth of opportunistic pathogens and such facilities should only use fresh inoculum.

The initial preservation trials also showed that glycerol was not an ideal cryopreservative in this system as it produced inoculum with lower overall survival, fewer viable bacterial species and an altered composition compared to the fresh starting material. This is of particular interest as glycerol is the standard cryopreservative used in microbiology laboratories^39^ and has been proposed for use in microbial biobanking settings. Our findings suggest that sucrose may prove to be a more successful cryopreservative for a greater range of microbial species and warrants further investigation and trial in the laboratory setting.

From the initial preservation trial, trehalose was eliminated as a potential excipient as it preserved fewer species compared to the other treatments. Further, the treatment where the three excipients (sucrose, trehalose and ascorbic acid) were used together did not provide increased survival over the single excipient treatments and therefore is not a favoured preservation method. While both ascorbic acid and sorbitol performed well in the initial preservation trials, in the longevity trial the viable cell composition was more dissimilar to the fresh material in the sorbitol treatment and ascorbic acid had lower richness. Interestingly, ascorbic acid had the highest viable cell richness in the initial preservation trials using wild koala scats. However, in the trial using scats from captive koalas, ascorbic acid favoured the survival of the family Muribaculaceae over other bacterial taxa. This suggests that performance of ascorbic acid as a preservative may be microbiome composition-dependent. Therefore, we concluded that the lyophilised fine particles without an added excipient or with the addition of sucrose are the best performing preservation methods in this study.

### Applications of koala faecal inoculation capsules

Our previous research has shown that koalas in different geographic regions have distinct gut microbiomes that are appropriate for digesting local *Eucalyptus* tree species^16^. Further, when koalas are translocated to a new habitat their gut microbiomes do not adapt to the change in diet but instead influence the *Eucalyptus* species they will feed on in the new habitat and their post translocation body condition^17^. Translocations have historically been used to manage koala populations in southern Australia where local densities have exceeded carrying capacity^40^. Translocations are also being increasingly used in northern Australian states to reestablish or supplement low density koala populations^41^. However, these translocations are often not successful with high mortality rates that in some cases have been linked to an inability of the koalas to feed on the local *Eucalyptus* foliage^40^. We have previously shown that our fresh faecal inoculation capsules can allow dietary expansion in captive koalas and there has been interest from state governments to use these capsules to pre-adapt the microbiomes of soon-to-be translocated koalas. Concurrently, captive ‘breed to release’ programs are being developed in both Queensland and NSW, where joeys of captive koalas will be used to boost wild population numbers. However, our research has demonstrated that the gut microbiomes of captive koalas are distinct from their wild counter parts^16,23^ and it is unknown if captively reared joeys will adapt to the diet in natural habitats. With their long shelf-life at room temperature, the lyophilised faecal inoculation capsules developed in this study could be readily used in a field setting to pre-adapt the microbiomes of koalas to the translocation site and to adjust the microbiomes of captively reared koalas pre-release.

Despite the successful development of a koala faecal inoculation with a prolonged shelf life, the capsules may not be readily adopted in a rehabilitation setting. In another unpublished study, we attempted to trial the use of the lyophilised faecal inoculation capsules in koalas receiving antibiotics with the aim of assessing whether they could prevent gut dysbiosis^5^. However, that trial was not successfully completed as veterinary and rehabilitation staff often struggled to reliably administer the capsules to the koalas. This led to increased real or perceived stress levels in the koalas receiving the capsules. Therefore, the provision of faecal inoculum via capsules may not be a viable option for the treatment of koalas due to varying expertise of the rehabilitation staff in capsule administration. As such, future research should focus on alternative delivery methods that can build upon the microbial preservation techniques developed and tested in this study. It is commonplace for koalas to be provided with nutritional supplements in the form of a paste administered by syringe (e.g. Crittacare Koala). The provision of the inoculum via this method may be better received by the sector. As such, the development of a microencapsulation method^18^ that can protect the preserved microbes in a powder formulation from koala stomach acid maybe offer a viable alternative.

An additional issue with koala faecal inoculation capsules that has become apparent during our research is the need for donor koalas. Firstly, it was very difficult to find appropriate donor koalas for our aborted *in vivo* study i) because of the need to use koalas with a “wild” faecal microbiome clashing with the practical necessity of using captive koalas to collect enough material to produce the capsules; and ii) because of the high prevalence of the exogenous koala retrovirus (KoRV) subtypes in northern koala populations that present a potential pathogen transmission risk^42–45^. Secondly, even where koalas with exogenous KoRV can be avoided, the use of donor koalas comes with an inherent risk of pathogen transmission. Thirdly, the time taken to collect and process the scats into fresh fine particles makes the use of donors impractical on a large scale. As such, for it to be feasible to produce inoculum for use in rehabilitation settings, it is necessary to find an alternative to donor koalas. One potential alternative would be the use of bioreactors to grow a microbial assemblage representative of a koala gut/faecal microbiome in the laboratory^46^.

## Conclusion

This study successfully developed a koala faecal inoculum with a shelf life of over a year that is viable for use in veterinary clinics and koala conservation programs such as translocations. However, in their current form the capsules are unlikely to be readily used in the rehabilitation setting due to challenges associated with administration and perceived stress to the koalas. The development of a powdered inoculum via microencapsulation that can be added to existing nutritional supplements may overcome this issue. Further, for the inoculum to be produced at scale and without the risk of pathogen transmission, further research into methods of growing a representative koala faecal microbiome community in the laboratory is needed.

## Methods

### Assessing initial survival and composition of faecal microbes post preservation

#### Starting material

In our first trial, we tested the initial survival of microbes derived from freshly collected scats from captive koalas housed at Lone Pine Koala Sanctuary. However, the high relative abundance of the bacterial family Muribaculaceae in those samples (typical of captive but not wild koala faecal microbiomes^16,23^) biased the overall survival in the different treatments towards excipients that favoured the survival of this family. As such, a second trial was conducted using scats from wild koalas excluding the air-dried treatments as those were found to be suboptimal (see Results).

In the second trial, initial survival and viable cell taxonomic composition of koala faecal microbes after preservation was assessed using scats collected from wild koalas at Belmont Hills Reserve, Brisbane, Australia. Fresh faecal pellets were collected from two koalas by placing a plastic mat (2 m by 3 m) directly beneath them late in the afternoon. Pellets were then collected from the mats the following morning and placed on ice until they could be returned to the laboratory for processing.

#### Preparation of formulations

The scats from the two koalas were pooled and then divided into 2-3 replicates of nine different treatments for preserving the faecal microbes as outlined in Table 1. Additionally, both fresh fine-particles and ground whole scats were assessed for comparison to the dried formulations. All treatments were prepared under anaerobic conditions in a Coy Vinyl Anaerobic Chamber containing a 95% nitrogen, 5% hydrogen atmosphere to maximise the survival of oxygen sensitive microbes. Fine particles were extracted from the scats as per the protocol of Blyton et. al. (2019). For the air-dried treatments, the scats or fine particles were left at room temperature for up to three days until dry. For the ground whole scat treatments, the scats were placed in an electric coffee grinder and ground until they were a fine powder. Five excipients were tested that have provided good survival of lyophilised bacteria over an extended period in other studies^19–22^ when used in the percentages listed in Table 1. The common excipients, skim milk powder and lactose, were not included in this study as koala milk contains low levels of lactose^24^ and adult koalas are thought to be lactose intolerant. Chosen excipients were added to the freshly prepared fine particles before lyophilisation. For the lyophilised treatments, the isolated fine particles were placed in a -80 °C freezer after addition of the relevant excipients for a minimum of 24 hours before being lyophilised in a BK-FD18 Biobase vertical freeze drier.

#### Assessment of survival

The proportion of viable microbes was determined using a LIVE/DEAD® BacLight™ Bacterial Viability Kit (Invitrogen, Thermo Fisher Scientific, Waltham, MA, USA, Cat. No. L7007) according to the manufacturer’s instructions. Briefly, a 1:100 dilution of 100 mg of sample was prepared using 1/4-strength Ringer’s solution (Merck Millipore, Ordering No. 1.15525.0001). Staining was performed by adding 0.54 µL of the BacLight dye mixture (equal parts Component A and B) to 180 µL of the dilution, followed by a 10-minute incubation in the dark. Stained samples (5µL) were then mounted onto 1% agarose slides. The proportion of viable, compromised, and dead cells were manually counted from images taken of the slides at 40x magnification using a ZEISS LSM 900 upright confocal microscope. Three technical replicates were imaged for each biological replicate per treatment. The BacLight Kit contains two stains. The first, SYTO9, passes through the cell membrane and stains DNA and RNA green, while the second, propidium iodine, stains DNA and RNA red but cannot penetrate the cell membrane^25^. Therefore, microbes with damaged cell membranes (presumed dead) are stained red using this protocol, while microbes with intact cell membranes (assumed viable) are stained green. We also identified a third category when applying this method to our samples where cells appeared yellow. These cells likely had compromised cell envelopes but not to the same extent as red cells.

### Assessment of viable cell taxonomic composition

To determine which microbial species survived, we extracted total DNA from each formulation after treatment with propidium monoazide (PMA), which binds to the DNA molecules of dead cells preventing their subsequent extraction^26^. After optimisation of the PMA treatment to our sample composition (see online supplementary), the addition of 0.25% 10mM PMA to a 1 in 2 dilution of the faecal material with a blue light exposure time of 30 minutes was found to effectively prevent extraction of DNA from dead cells without causing additional cell death and was subsequently used in all experiments.

For each sample, total DNA was extracted from approximately 50mg of material. The material was beaten for 10 minutes using the MoBio PowerLyzer24 in a MoBio bead tube containing 0.1 mm diam. Zirconian/silica beads and 750ul of TLA buffer (Promega). The samples were centrifuged at 10,000 g for 30 seconds. DNA was then extracted from 200 ul of the supernatant using the Maxwell 16 robotic system and corresponding Tissue DNA kit (Promega) following the manufacturer’s instructions.

16S rRNA amplicon sequencing was then performed using the workflow outlined by Illumina (#15044223 Rev.B). A 589 bp section of the 16S rRNA gene (V5 – V8 region) was amplified using 803F and 1392R primers^27^ with the addition of Illumina specific adapter sequences, according to the specified workflow except that NEBNext® Ultra™ II Q5® Mastermix (New England Biolabs #M0544) was used in place of the standard workflow polymerase. PCR amplicons were purified using Agencourt AMPure XP beads (Beckman Coulter). Purified products were indexed using the Illumina Nextera XT 384 sample Index Kit A-D (Illumina FC-131-1002). Indexed amplicons were pooled in equimolar concentrations and sequenced on a MiSeq (Illumina) using paired end sequencing with V3 2x300bp chemistry at the Australian Centre for Ecogenomics.

Primer sequences were removed from forward de-multiplexed sequencing reads using cutadapt (version 2.10)^28^, with reads not containing primers discarded. Poor quality reads were identified and removed with trimmomatic (version 0.39)^29^ using a sliding window of 4 bases with an average quality of 15. Reads were trimmed to 250bp, with those less than 250bp discarded. Quality controlled forward reads were then processed using QIIME2 (ver. 2020.11.1)^30^ for feature selection, abundance calculations and taxonomy assignment. Reads were de-noised using DADA2^31^ and the relative abundance of each resulting feature calculated. The taxonomy was assigned using the combined non-redundant 16S and 18S SILVA database (release 138, clustered at 99% identity)^32^ by BLAST using the classify-consensus-blast function with default parameters. The resulting microbial feature-by-sample tables were rarefied to 9000 reads per sample using the vegan package^33^ in R^34^. All community composition analyses were performed on the rarefied dataset. Microbiome richness was estimated by calculating the Chao Index of alpha diversity using the package fossil in R^35^. Dissimilarity between samples was estimated by Bray-Curtis distances calculated using the vegan package.

### Assessing the survival and composition of preserved and encapsulated faecal microbes over a year

Scats from three wild born koalas housed at Currumbin Wildlife Hospital were used as the starting material for the long-term preservation trial. To obtain enough material for the experiment, scats were collected daily from the base of the koalas’ enclosures over a period of three days, with the fine particles extracted, excipients added and then frozen at -80°C each day as above. After collections were complete the processed material was lyophilised as above and pooled across the three days and koalas for each treatment. Treatments 5, 6, 7 and 11 (Table 1) were selected for this trial (see results) and the material prepared for each as above. The material for each treatment was then packaged into two hypromellose capsules (DRCaps, Capsugel®) and coated in shellac solution (37% w/v shellac, 61.5% v/v ethanol and 1.5% v/v Tween 20). Once the shellac solution had dried, the capsules were placed in amber airtight glass jars with at least three desiccant pouches (MiniPax). The jars were kept at room temperature in the laboratory (21 – 26°C) for up to a year.

Each treatment was assessed at 13 timepoints (Table 2) for overall microbial survival and viable cell taxonomic microbial composition. At each time point, three capsules from each treatment were removed from the amber jars and cut open to access the material inside. After the material was reconstituted in ¼ Ringers solution it was stained with SYTO9 and propidium iodine^25^ and the proportion of viable, compromised, and dead cells determined from cell counts using confocal microscopy as described above. To determined which microbial species survived, total DNA was extracted from the material after treatment with propidium monoazide (PMA), 16S rRNA gene amplicon sequencing performed and bioinformatic analysis conducted as above. Seven samples including three fresh fine particle replicates and a single time zero replicate sample were included in each sequencing run so that inter-run differences could be assessed and accounted for.

**Table 2:** Long term preservation study sampling timepoints.

| Time point | Weeks into trial (Days) |
| --- | --- |
| 0 | 0 (0) |
| 1 | 1 (7) |
| 2 | 2 (14) |
| 3 | 3 (21) |
| 4 | 5 (35) |
| 5 | 7 (49) |
| 6 | 9 (63) |
| 7 | 11 (77) |
| 8 | 15 (105) |
| 9 | 23 (161) |
| 10 | 31 (217) |
| 11 | 41 (287) |
| 12 | 52 (365) |

To assess changes in the capsules over time, linear regression models were constructed in R for Chao1 richness and rank transformed Bray-Curtis distance to the time zero replicate from the relevant treatment included on the same sequencing run. The number of days into the longevity trial and the treatment were fitted as explanatory variables with an interaction term considered. The sequencing run was also included as a covariate in the analyses to account for inter-run differences. The assumption that the residuals of the model conformed to normality was confirmed by Shapiro-Wilkes test.

For visualisation purposes, Chao1 richness and Bray-Curtis distances were adjusted for all samples in a sequencing run to account for inter-run differences. For Chao1, this was done by subtracting the average Chao1 richness for the seven repeat samples on the run [Chao1_rr_] from a sample’s Chao1 richness [Chao1_S_] and then adding the average Chao1 richness for the repeat samples across all runs [Chao1_ar_] (Chao1_S_ – Chao1_rr_ + Chao1_ar_). For Bray-Curtis distances to the T0 samples this was done by subtracting the average Bray-Curtis distance among T0 replicates on the run [Bray_T0rr_] from the Bray-Curtis distance to T0 for the sample [Bray_s_] and then adding the average Bray-Curtis distance among T0 replicates across all runs [Bray_T0ar_] (Bray_s_ - Bray_T0rr_ + Bray_T0ar_).

### Determination of optimal capsule design for dry fill

Our previous research demonstrated that the wet fill capsule design was effective at delivering viable microbes past the acid stomach to the koala hindgut^14^. In that experiment, we first tested the capsule design *in vitro* and showed that two hypromellose capsule (DRCaps, Capsugel®) layers coated in a shellac solution (37% w/v shellac, 61.5% v/v ethanol and 1.5% v/v Tween 20) was necessary for the capsule to maintain integrity for 10 hours in an acidic solution (replicating transit time in a koala stomach). By contrast, single layered capsules quickly lost integrity due to the internal moisture of the inoculum. However, in this study, we packaged dry material into the capsules. Thus, we repeated our *in vitro* experiment with dry filled capsules to determine if single layer capsules could be used, enabling a larger volume of inoculum to be administered in a single capsule. The experiment was completed as per Blyton et al.^14^ except that the capsules were filled with lyophilised fine particles with 16.7% sorbitol produced in the long-term preservation trial and mixed with a small quantity of crystal violet powder.

## Acknowledgements

We thank Sean FitzGibbon who facilitated our collection of koala scats from the wild koalas in Belmont Hills Reserve and Mariska de Bruin for her assistance in preparing the lyophilised faecal capsules for the longevity study. We also thank Daisy Hill Koala Centre, Lone pine Sanctuary and Currumbin Wildlife Hospital for providing us with the faecal material from the donor koalas for our experiments. We acknowledge the traditional owners of the lands on which this work was conducted, the Jagera, Turrbal and Yugambeh peoples. Samples were collected under University of Queensland ethics approval (2022_AE000533). This project was funded by the Brisbane City Council, Australia, as part of their Koala Research Partnerships Program. The funders had no role in study design, data collection and interpretation, or the decision to submit the work for publication.

